# Perfect 21-nucleotide matches to beneficial fungi are common in canonical antifungal dsRNA targets: an in-silico off-target hazard screen for spray-induced gene silencing

**DOI:** 10.64898/2026.09.22.753319

**Authors:** Muhammad Wasif, Gábor Tarcali, Muhammad Ahsan Khan, Huma Saleem, Muhammad Abubakkar Azmat

**Author notes:** Corresponding author: Muhammad Wasif, Institute of Plant Protection, Faculty of Agricultural and Food Sciences and Environmental Management, University of Debrecen, Böszörményi út 138, 4032 Debrecen, Hungary. Co-author e-mails: Gábor Tarcali; Muhammad Ahsan Khan; Huma Saleem; Muhammad Abubakkar Azmat.

## Abstract

Double-stranded RNA (dsRNA) biopesticides that silence essential fungal genes by spray-induced gene silencing (SIGS) are advancing towards registration, underpinned by the claim that silencing is confined to sequence-matched target organisms, a claim that has never been tested systematically against beneficial fungi. We screened twelve dsRNA constructs (the whole transcripts of eleven canonical antifungal target genes of *Fusarium graminearum*: three CYP51 sterol 14α-demethylases, six chitin synthases and two β-tubulins, plus a concatenation of the three CYP51 transcripts modelling the flagship CYP3RNA design) for perfect 21-nucleotide (21-nt) identity, the primary match criterion of recent regulatory bioinformatics frameworks, against the transcriptomes of seven non-target fungi, including commercial biocontrol agents (*Trichoderma harzianum*, *T. virens*, *Beauveria bassiana*, *Metarhizium robertsii*, *M. brunneum*), the arbuscular mycorrhizal fungus *Rhizophagus irregularis* and *Saccharomyces cerevisiae*. Eleven of the twelve constructs carried off-target hazard; only CYP51C was completely clean. TUB2b carried 580 perfect 21-nt matches and contained no hazard-free window of 250 consecutive 21-mer start positions, whereas 26.6–60.0% of screened windows were hazard-free in the designable genes. Hits landed overwhelmingly on orthologues of the targeted gene and concentrated in the Hypocreales; *R. irregularis* and *S. cerevisiae* were nearly clean. These are hazard findings, not risk findings: perfect siRNA-length matches to beneficial fungi are common in canonical antifungal dsRNA targets, and current regulatory screening panels contain no fungus that would detect them.

## Introduction

RNA interference (RNAi) has moved from a laboratory tool to a crop-protection technology. Host-induced gene silencing (HIGS), in which a transgenic plant expresses RNAi constructs against transcripts of an invading pathogen, was first demonstrated against the powdery mildew fungus *Blumeria graminis* in barley and Arabidopsis (Nowara et al. 2010), and was extended to *Fusarium graminearum* by Koch et al. (2013), who showed that silencing the three fungal CYP51 sterol 14α-demethylase genes (the targets of azole fungicides) conferred strong resistance. Spray-induced gene silencing (SIGS) removed the need for transgenesis: Wang et al. (2016) showed that *Botrytis cinerea* takes up environmental RNAs and that spraying Dicer-gene-targeting dsRNAs controlled grey mould on fruits, vegetables and flowers, while Koch et al. (2016) demonstrated that a sprayed ∼800-nt dsRNA (CYP3RNA) complementary to all three *FgCYP51* paralogues reduced *F. graminearum* infection of barley in a process dependent on the fungal silencing machinery. Subsequent work compared the two delivery strategies directly (Koch et al. 2019), dissected how construct length shapes the siRNA pool and silencing efficiency (Höfle et al. 2020), and developed formulations such as layered double-hydroxide clay nanosheets that extend the persistence of sprayed dsRNA (Mitter et al. 2017; Sede et al. 2026).

Commercially, RNAi crop protection has so far been realised against animal pests. Transgenic maize expressing DvSnf7 dsRNA against western corn rootworm completed ecological risk assessment on the strength of bioinformatic match analysis combined with tiered non-target bioassays (Bachman et al. 2016), and in December 2023 the United States Environmental Protection Agency registered ledprona (product Calantha), the first sprayable dsRNA biopesticide, targeting the Colorado potato beetle proteasome subunit gene *PSMB5* (US EPA 2023). A second registered dsRNA product, the anti-*Varroa* acaricide vadescana (marketed as Norroa and registered by the US EPA in September 2025), has followed the same regulatory path, although it is applied inside the hive rather than sprayed onto a crop (Lester et al. 2026). No antifungal SIGS product has yet reached registration, but the antifungal pipeline is scientifically the most mature of the remaining candidates (Koch et al. 2019; Dietz-Pfeilstetter et al. 2021), which makes the environmental safety questions specific to antifungal constructs timely rather than hypothetical.

The central safety claim of the dsRNA biopesticide field is sequence specificity: because silencing is triggered by base-pairing between dsRNA-derived siRNAs and complementary mRNA, effects are expected to be confined to organisms that carry near-identical target sequence. This claim was imported from the insecticidal literature, where it is empirically well supported. Baum et al. (2007) noted that insecticidal activity tracked matching sequence in the target pest, and Bachman et al. (2013) systematically characterised the activity spectrum of the DvSnf7 dsRNA across insects, finding activity essentially confined to close relatives of the western corn rootworm, which established the rule that off-target activity scales with phylogenetic relatedness and sequence identity. Within that literature the rule is reassuring, because the beneficial organisms of regulatory concern (pollinators, predators, aquatic arthropods) are phylogenetically distant from the target pests.

Antifungal dsRNA constructs break the premise of that reassurance. The beneficial fungi that agriculture actually deploys are not distant relatives of the target pathogens; they are their phylogenetic neighbours. *Fusarium*, the genus against which the flagship antifungal constructs are designed, sits in the order Hypocreales alongside *Trichoderma* spp., the most widely commercialised fungal biocontrol agents (Harman et al. 2004; Lorito et al. 2010), and alongside the entomopathogenic genera *Beauveria* and *Metarhizium* that underpin the mycoinsecticide industry (Mascarin and Jaronski 2016; St. Leger and Wang 2020). The canonical antifungal target genes (sterol demethylases, chitin synthases, β-tubulins) are essential, conserved housekeeping genes whose orthologues are present in every filamentous fungus. The relatedness–spectrum rule of Bachman et al. (2013) therefore predicts, quantitatively and uncomfortably, that the off-target sequence space of an antifungal dsRNA should be enriched precisely in the beneficial fungi most likely to be co-deployed with it. Whether this prediction holds has, to our knowledge, never been tested systematically.

The operational tool for such a test already exists. Regulatory bioinformatics for dsRNA biopesticides has converged on a sequence-based hazard criterion: any contiguous stretch of at least 21 nucleotides of perfect identity between the dsRNA and a non-target transcript (the length of a Dicer-processed siRNA) is flagged as a potential silencing trigger warranting further scrutiny, complemented by overall-identity thresholds of approximately 80% (De Neef et al. 2026; Devisetty et al. 2025; both frameworks proposed by GreenLight Biosciences), an approach also reflected in OECD considerations for sprayed and externally applied dsRNA-based pesticides (OECD 2020, 2023). The approach has precedent in pollinator safety research: k-mer matching showed that pesticidal dsRNAs can carry short perfect matches to the honey bee genome (Mogren and Lundgren 2017), though experimental follow-up is essential: in the bumble bee *Bombus terrestris* (Taning et al. 2021), none of 24 transcripts predicted as off-targets by sequence complementarity, including one carrying a 20-nt perfect match, showed significant knockdown after ingestion of the dsRNA, underscoring that a sequence match flags hazard rather than demonstrating effect. The criterion is deliberately conservative at the sequence level and is understood to flag hazard, not effect: mismatch-tolerant silencing below the threshold is also documented (Jackson et al. 2003; Kulkarni et al. 2006; Chen et al. 2021), so a perfect-match screen bounds only the most obvious component of off-target sequence space.

The criterion has particular mechanistic force in fungi. Most fungi possess a canonical RNAi pathway in which Dicer-like RNase III enzymes process long dsRNA into siRNAs of roughly 21–25 nt that load onto Argonaute proteins and direct cleavage of complementary transcripts, and because Argonaute-mediated slicing requires extensive complementarity along the siRNA duplex, a contiguous perfect ∼21-nt match is the biologically meaningful unit of off-target concern (Torres-Martínez and Ruiz-Vázquez 2017). Fungal systems also demonstrate that complementarity-driven cross-reactivity is efficient: a single dsRNA silences all three CYP51 paralogues of *F. graminearum* simultaneously through the identical stretches they share (Koch et al. 2016), and longer constructs generate more diverse siRNA pools with correspondingly greater silencing and cross-silencing potential (Höfle et al. 2020). Finally, the exposure route is real: pathogenic fungi take up RNA from their environment, and this uptake is the mechanistic basis of SIGS itself (Wang et al. 2016; Qiao et al. 2021). The same machinery that makes antifungal dsRNA sprays work is the machinery through which an off-target match in a non-target fungus would act.

There is, however, a structural gap in where these screens are pointed. Non-target organism selection in current ecological risk assessment is exposure-driven and comprises guideline indicator species (bees, arthropods, earthworms, aquatic taxa, vertebrates) but no fungus (Romeis and Widmer 2020; Dietz-Pfeilstetter et al. 2021). Under Regulation (EC) No 1107/2009, plant protection products must show no unacceptable effects on the environment, and the EFSA Panel on Plant Protection Products and their Residues has framed specific protection goals for in-soil organisms around ecosystem services (EFSA PPR Panel 2017); and EFSA’s 2024 materials for revising the terrestrial ecotoxicology guidance identify tests on soil fungi as a gap not covered by the current test battery (EFSA 2024). Arbuscular mycorrhizal fungi (AMF) have been proposed for inclusion in the test battery (Mallmann et al. 2018), and method development for AMF ecotoxicity testing has frequently focused on *Rhizophagus irregularis* as the model species for the soil-microorganism protection goal (Roshanfekrrad et al. 2025; Papadopoulos et al. 2025), proposed but not yet mandatory. Fungal biocontrol agents are themselves approved active substances at strain level, but a commercial *Trichoderma* strain is protected when it is the product and unprotected as a non-target organism when another product is assessed.

This paper reports a systematic in-silico off-target hazard screen of canonical antifungal dsRNA targets against beneficial and non-target fungi. The question is deliberately narrow and tractable: if a dsRNA is designed anywhere within a canonical antifungal target gene of *F. graminearum*, how often does it contain a stretch of 21 nucleotides that perfectly matches a transcript in a beneficial or non-target fungus? Twelve constructs (eleven whole target transcripts spanning the CYP51, chitin synthase and β-tubulin families, plus a concatenation of the three CYP51 transcripts, CYP51-concat, modelling the CYP3RNA design) were screened against the transcriptomes of seven non-target fungi representing the major guilds of deployed beneficial fungi. Throughout, the analysis is framed as a hazard screen, not a risk assessment: hazard asks whether a silencing trigger is present in the sequence, whereas risk additionally requires environmental exposure, uptake by the non-target organism, and a functioning silencing pathway (Qiao et al. 2021; De Neef et al. 2026). The remainder of the paper describes the reference panel, query set and exact-match computational method (Section 2); reports construct-level hit counts, the identity of matched transcripts, the phylogenetic distribution of hazard, and a sliding-window designability analysis (Section 3); and discusses the interpretation, limitations and regulatory implications of the findings (Section 4).

## Materials and methods

### Reference panel

Reference transcriptomes (cDNA) were downloaded from Ensembl Fungi on 3 August 2026 (Yates et al. 2022; Dyer et al. 2025); Ensembl Fungi release 63. The panel comprised twelve species, 158,032 transcripts and 218 Mb of sequence (Table 1). Species were chosen to span the functional guilds of fungi relevant to an antifungal dsRNA spray scenario rather than to sample fungal diversity evenly. *Fusarium graminearum* PH-1 is the target pathogen. Seven species constitute the non-target panel: the commercial biocontrol agents *Trichoderma harzianum* and *T. virens* (Harman et al. 2004); the entomopathogens *Beauveria bassiana*, *Metarhizium robertsii*, and *M. brunneum*, represented by its sequenced reference-genome strain ARSEF 3297, which is not itself a commercial isolate (the deployed biocontrol strains of this species are others, notably F52/BIPESCO 5, approved as active substances in the EU and US), chosen so that a hit falls on the genome of the commercially deployed entomopathogenic genus *Metarhizium* biocontrol agents (strain-level *Metarhizium* products are approved as active substances in the EU and US), so that a hit there is a hit on a commercially deployed class of biocontrol agent (Mascarin and Jaronski 2016; St. Leger and Wang 2020); the arbuscular mycorrhizal fungus *Rhizophagus irregularis* DAOM 181602, chosen deliberately because it is the species around which AMF test-method development for the soil-microorganism protection goal has largely centred (Roshanfekrrad et al. 2025; Papadopoulos et al. 2025); and *Saccharomyces cerevisiae* as a well-characterised reference fungus that additionally serves as a natural negative control, because it lacks functional Dicer and Argonaute genes and therefore cannot mount canonical RNAi (Drinnenberg et al. 2009). Four further plant pathogens (*Botrytis cinerea*, *Sclerotinia sclerotiorum*, *Verticillium dahliae*, *Zymoseptoria tritici*) were included as comparison species for future expansion of the screen; the present analysis reports non-target results only.

**Table 1.** Reference panel. Transcriptomes (cDNA) were retrieved from Ensembl Fungi on 3 August 2026 (release 63; 12 species, 158,032 transcripts, 218 Mb). The seven non-target species screened in this study are listed between the target pathogen and the comparison pathogens. The strain ARSEF 23, whose genome was originally deposited as Metarhizium anisopliae, has since been reclassified as M. robertsii (Bischoff et al. 2009).

| Role | Species | Assembly |
| --- | --- | --- |
| Target pathogen | *Fusarium graminearum* PH-1 | ASM24013v3 |
| Beneficial, biocontrol | *Trichoderma harzianum* | ASM98886v1 |
| Beneficial, biocontrol | *Trichoderma virens* | ASM17099v1 |
| Beneficial, entomopathogen | *Beauveria bassiana* | ASM168263v1 |
| Beneficial, entomopathogen | *Metarhizium robertsii* ARSEF 23 | MAA_2.0 |
| Beneficial, entomopathogen | *Metarhizium brunneum* ARSEF 3297 | MBR_1.0 |
| Beneficial, arbuscular mycorrhizal | *Rhizophagus irregularis* DAOM 181602 | ASM43914v3 |
| Reference fungus | *Saccharomyces cerevisiae* | R64-1-1 |
| Comparison pathogen | *Botrytis cinerea* | ASM83294v1 |
| Comparison pathogen | *Sclerotinia sclerotiorum* | ASM14694v1 |
| Comparison pathogen | *Verticillium dahliae* | ASM15067v2 |
| Comparison pathogen | *Zymoseptoria tritici* | MG2 |

### Query set

Eleven canonical antifungal dsRNA target genes were taken from the *F. graminearum* annotation: the three sterol 14α-demethylase genes CYP51A, CYP51B and CYP51C (locus tags FGSG_04092, FGSG_01000 and FGSG_11024), which are the genes targeted by CYP3RNA, the most-cited antifungal dsRNA construct in existence (Koch et al. 2013, 2016); six chitin synthases (CHS1, FGSG_10116; CHS2, FGSG_02483; CHS3, FGSG_10327; CHS4, FGSG_01272; CHS6, FGSG_12039; CHSD, FGSG_01949); and two β-tubulins (TUB2a, FGSG_06611, the β2-tubulin, and TUB2b, FGSG_09530, the β1-tubulin; Zhu et al. 2021), selected as canonical antifungal RNAi targets from the *F. graminearum* annotation. A concatenation of all three CYP51 transcripts was additionally screened as a twelfth construct, CYP51-concat, modelling the design logic of CYP3RNA. Queries were whole target transcripts, not the exact published construct fragments; the numbers therefore describe the hazard of designing a dsRNA anywhere within each gene, rather than the hazard of any specific published sequence. Screening the actual published construct sequences, the corpus audit, is identified as future work.

### The 21-nt perfect-identity criterion

A hit was defined as a contiguous stretch of 21 nucleotides of perfect identity between the construct and a non-target transcript. Twenty-one nucleotides is the length of a canonical Dicer-processed siRNA in fungi, where Dicer-like RNase III enzymes process long dsRNA into small RNAs of approximately 21–25 nt that load onto Argonaute and direct cleavage of complementary transcripts (Torres-Martínez and Ruiz-Vázquez 2017). It is also the primary match criterion recommended for regulatory off-target screening of externally applied dsRNA biopesticides, where a ≥21-nt perfect-identity stretch against a non-target transcriptome is treated as a hazard flag requiring further scrutiny (De Neef et al. 2026), a criterion with operational precedent in pollinator hazard screening (Mogren and Lundgren 2017; Taning et al. 2021). The second component of the recommended criterion, ∼80% overall sequence identity, requires alignment and was not implemented here (see Sections 2.4 and 4). A perfect-match criterion also has known limits: mismatch-tolerant off-target silencing by short duplexes is well documented experimentally (Jackson et al. 2003; Kulkarni et al. 2006; Chen et al. 2021), so the screen bounds only the most stringent component of off-target sequence space (Section 4).

### Exact-match computation

The screen uses exact k-mer membership testing rather than alignment, in four steps.

*Step 1: two-bit encoding.* Every 21-mer was encoded as a single 64-bit integer by mapping A, C, G, T to 0, 1, 2, 3. A 21-mer becomes a number below 4^21 (≈ 4.4 × 10^12), which fits comfortably in a 64-bit integer. Encoding used Horner’s method (start at zero, and for each of the 21 positions multiply the running value by four and add the base value), vectorised in numpy as 21 array passes rather than a Python loop over millions of k-mers. Comparing integers is far faster and uses far less memory than comparing strings: a whole fungal transcriptome reduces to a sorted array of roughly 20 million integers (∼150 MB), which is why the analysis runs on a laptop instead of a cluster.

*Step 2: ambiguity filtering.* Any 21-mer window overlapping an ambiguous base (N or other IUPAC ambiguity codes) was discarded, because perfect identity cannot be claimed through an unresolved position. The filter was implemented with a cumulative sum over an ambiguity mask so that it remains vectorised.

*Step 3: both-strand querying.* A dsRNA duplex is diced into siRNAs derived from both strands, so a match in either orientation is a potential silencing trigger. The query set was therefore the union of the 21-mers of each construct and of its reverse complement, queried against an index built from the sense strand of reference transcripts only. This design has an explicit built-in positive control: exactly half of the query k-mers (the sense half) can match a sense-strand index, so the self-hit rate of *F. graminearum* against its own genes is expected to be ≈50%; in the run reported here it read exactly 50.00%, confirming that the pipeline behaved as intended, and any drift from 50% would indicate a defect (Section 3.1).

*Step 4: lookup, not alignment.* Membership testing used numpy.searchsorted against the sorted reference array. The lookup is exact and deterministic, with none of the heuristic seeding used by aligners such as BLAST or bowtie; for a perfect-identity criterion this is both faster and more correct than an aligner, and it complements design-oriented tools such as si-Fi and integrated design platforms that couple siRNA prediction with off-target search (Lück et al. 2019; Cedden et al. 2025).

One construct-specific check applies to CYP51-concat: concatenating the three CYP51 transcripts creates two artificial junctions, each generating 20 junction-spanning 21-mers (40 in total) that do not exist in any natural transcript; these were checked explicitly and contributed zero hits: the per-species hit counts of the concatenation equal the exact sum of the three component genes (Table 2).

**Table 2.** Off-target hazard across constructs. Number of distinct 21-mers of each construct (either strand) with a perfect match in each non-target transcriptome. len, construct length in nucleotides; total, hits summed across the seven non-target species.

| Construct | len<br>nt | *T.*<br>*harz.* | *T.*<br>*vir.* | *B.*<br>*bass.* | *M.*<br>*rob.* | *M.*<br>*brun.* | *R.*<br>*irreg.* | *S.*<br>*cer.* | total |
| --- | --- | --- | --- | --- | --- | --- | --- | --- | --- |
| CYP51-concat | 4659 | 35 | 10 | 37 | 21 | 26 | 0 | 0 | 129 |
| CYP51A | 1524 | 0 | 0 | 3 | 0 | 1 | 0 | 0 | 4 |
| CYP51B | 1581 | 35 | 10 | 34 | 21 | 25 | 0 | 0 | 125 |
| CYP51C | 1554 | 0 | 0 | 0 | 0 | 0 | 0 | 0 | 0 |
| CHS1 | 3408 | 112 | 156 | 15 | 31 | 31 | 3 | 0 | 348 |
| CHS2 | 2478 | 6 | 6 | 18 | 0 | 0 | 0 | 0 | 30 |
| CHS3 | 2879 | 24 | 16 | 0 | 5 | 12 | 0 | 0 | 57 |
| CHS4 | 3573 | 8 | 0 | 23 | 9 | 9 | 1 | 0 | 50 |
| CHS6 | 4677 | 28 | 16 | 31 | 1 | 1 | 0 | 0 | 77 |
| CHSD | 2328 | 3 | 0 | 3 | 0 | 0 | 1 | 0 | 7 |
| TUB2a | 1686 | 37 | 46 | 22 | 3 | 3 | 10 | 3 | 124 |
| TUB2b | 1648 | 120 | 133 | 80 | 118 | 126 | 3 | 0 | 580 |

### Designability analysis

To ask whether observed hazard is avoidable by construct placement, a sliding-window analysis was performed in k-mer coordinates. A window of 250 consecutive 21-mer start positions, spanning 270 nt of transcript because a window of n start positions covers n + 20 nucleotides, was slid along each gene in steps of one start position, and each window was classified as clean if it contained zero hazardous 21-mers against any of the seven non-target fungi. For each gene the analysis reports the total number of windows, the number and percentage of clean windows, and the worst window (the maximum number of hazardous 21-mers in any single window). The positional distribution of hazardous 21-mers along the five screened transcripts (CYP51A, CYP51B, CYP51C, CHS1 and TUB2b) was visualised as per-position hazard landscapes (Figure 1).

**Fig. 1.**
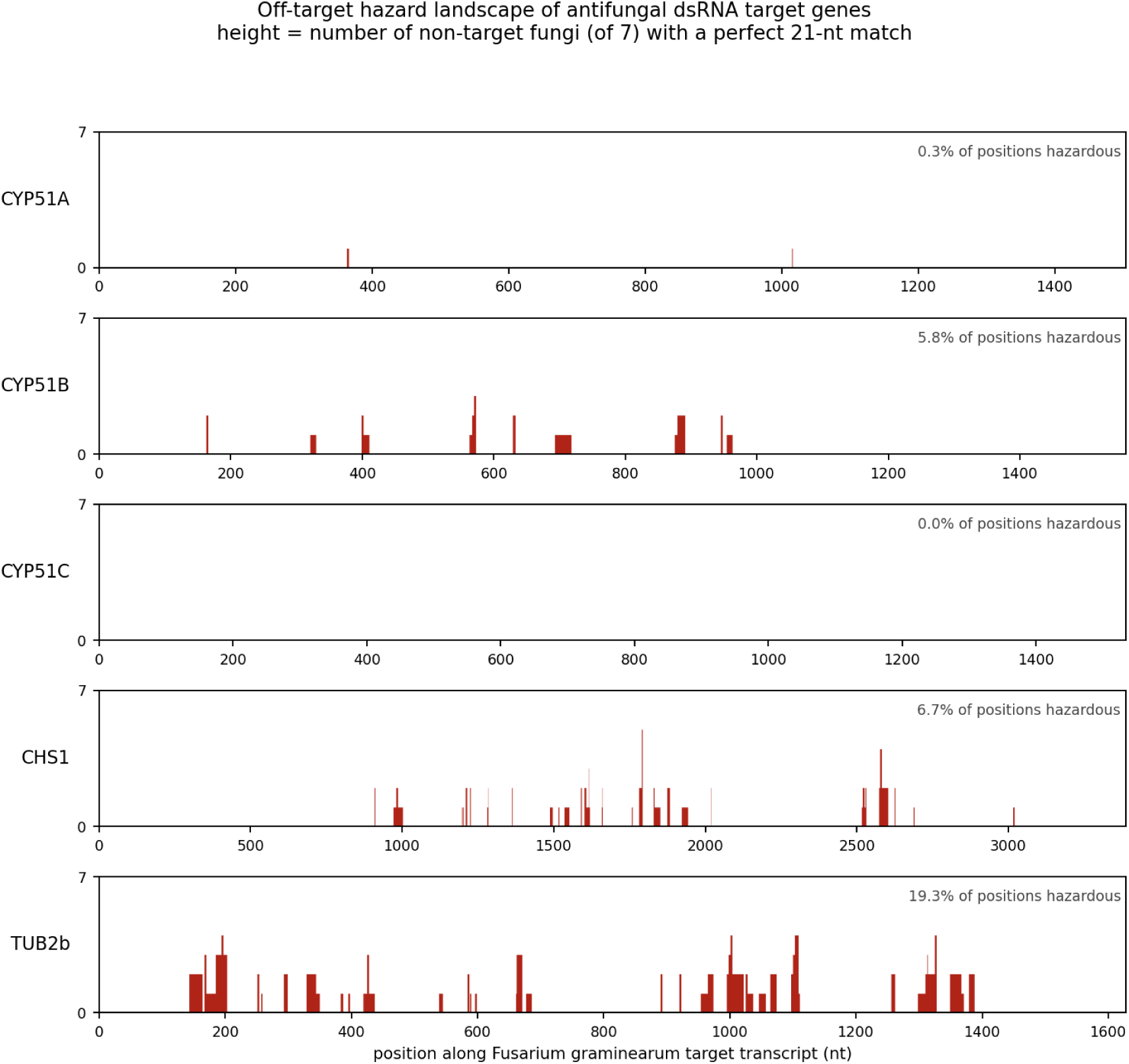
Positional off-target hazard landscapes for five canonical antifungal dsRNA target genes of *Fusarium graminearum* (CYP51A, CYP51B, CYP51C, CHS1, TUB2b). Each panel shows, for every position along the target transcript, the number of the seven screened non-target fungi (*Trichoderma harzianum*, *T. virens*, *Beauveria bassiana*, *Metarhizium robertsii*, *M. brunneum*, *Rhizophagus irregularis*, *Saccharomyces cerevisiae*) whose transcriptomes contain a perfect 21-nt match to the 21-mer beginning at that position; height therefore ranges from 0 (no non-target match) to 7 (a match in every non-target species). The percentage annotated per panel is the density of hazardous start positions, i.e. the fraction of transcript positions whose 21-mer has at least one perfect non-target match (e.g., CYP51A: 4 hazardous 21-mers over ∼1,500 possible start positions ≈ 0.3%), not the fraction of transcript covered by hazardous 21-mers: CYP51A, 0.3%; CYP51B, 5.8%; CYP51C, 0.0%; CHS1, 6.7%; TUB2b, 19.3%. Hazard is concentrated in discrete peaks corresponding to conserved blocks of orthologous essential genes, most densely in TUB2b, consistent with the sliding-window designability analysis in Table 3

**Table 3.** Designability analysis. A window of 250 consecutive 21-mer start positions (spanning 270 nt of transcript) was slid along each gene in steps of one start position; a window is clean if it contains no 21-mer with a perfect match in any of the seven non-target fungi. Worst window, the maximum number of hazardous 21-mers in any single window.

| Gene | windows | clean windows | clean % | worst window |
| --- | --- | --- | --- | --- |
| CYP51C | 1285 | 1285 | 100.0 | 0 |
| CYP51A | 1255 | 753 | 60.0 | 3 |
| CHS1 | 3139 | 1102 | 35.1 | 67 |
| CYP51B | 1312 | 349 | 26.6 | 41 |
| TUB2b | 1379 | 0 | 0.0 | 100 |

### Reproducibility

The complete analysis is reproducible from public data on a laptop, with no wet laboratory, no compute cluster and no paid software. Analyses were run in Python 3 using numpy, Biopython and matplotlib; the analysis code and result files are archived at Harvard Dataverse (see Data availability). The engine (kmerlib.py) is 86 lines of commented Python built on numpy and Biopython; companion scripts download the reference panel, run the screen in batches of two species per call, and generate the positional hazard maps. Downloading the full reference panel (218 Mb) completes within minutes over a standard broadband connection. The run reported here was executed on 3 August 2026.

## Results

### Eleven of twelve constructs carry off-target hazard

The built-in positive control validated the pipeline before any non-target result was read: screening *F. graminearum* against its own transcriptome returned a self-hit rate of exactly 50.00%, the value predicted by the both-strand query design (Section 2.4). Every 21-mer of each of the twelve constructs (and of its reverse complement) was then tested for perfect identity against the seven non-target fungal transcriptomes. Table 2 reports the number of 21-mers with at least one perfect match in each non-target species. The screen found off-target sequence hazard to be the rule rather than the exception: eleven of the twelve constructs carried at least one perfect 21-nt match to a non-target fungus. Only CYP51C was completely clean, with zero matches in all seven species. Hazard levels spanned more than two orders of magnitude, from four hits in CYP51A to 580 in TUB2b. The β-tubulins and the chitin synthase CHS1 were the heaviest carriers (TUB2b, 580 hits; CHS1, 348; TUB2a, 124), while the CYP51 family was sharply internally divided: CYP51B carried 125 hits, CYP51A carried four, and CYP51C none. The CYP51-concat concatenation carried 129 hits, of which 125 (97%) were attributable to the CYP51B component; that is, essentially all of the off-target sequence hazard in a construct modelled on the field’s flagship design is concentrated in one of its three gene components.

### Hits land on orthologues of the targeted gene

Where the hits land matters more than the raw counts. Inspection of the matched non-target transcripts showed that the matches are not random sequence: they are the orthologues of the targeted gene. TUB2b 21-mers matched β-tubulin transcripts in *T. harzianum*, *M. brunneum* and *R. irregularis*; CYP51B 21-mers matched eburicol 14-α-demethylase (the CYP51 orthologue) in *T. harzianum* and cytochrome P450 51B in *M. brunneum*; and CHS1 21-mers matched chitin synthase 1 in *T. harzianum* and *M. brunneum*. In other words, a dsRNA designed to kill a pathogen by silencing an essential gene carries perfect siRNA-length matches to the same essential gene in commercially deployed biocontrol fungi. One annotation caveat applies: several hits in *B. bassiana* and *T. virens* land on transcripts annotated only as “hypothetical protein”, which are almost certainly the unannotated orthologues but cannot be stated as such without further checking (Section 4).

### Hazard tracks phylogeny

The distribution of hits across species followed phylogeny rather than transcriptome size or assembly quality. The five non-target Hypocreales (*Trichoderma*, *Beauveria* and *Metarhizium*, the same order as the target genus *Fusarium*) carried the overwhelming majority of hits in every construct (Table 2): 1,510 of the 1,531 hazardous 21-mers (98.6%) occurred in a Hypocreales non-target. The two phylogenetically distant species were nearly clean: *R. irregularis* (Glomeromycota) carried at most ten hits against any construct (TUB2a), and zero against the entire CYP51-concat concatenation, while *S. cerevisiae* carried only three hits in total, all against TUB2a. This is the quantitative version of the relatedness–spectrum argument: the beneficial fungi that agriculture deploys against or alongside *Fusarium* are its phylogenetic nearest neighbours, and the specificity reasoning imported from the insecticidal dsRNA literature, where a honey bee is not a close relative of a corn rootworm in the way *Trichoderma* is a close relative of *Fusarium*, does not transfer (Bachman et al. 2013). Two control observations anchor the interpretation. First, the near-clean result for *R. irregularis* is good news for mycorrhizae and is reported as such: the species around which AMF regulatory test development has converged carries negligible perfect-match hazard from these constructs. Second, the near-zero count in *S. cerevisiae* behaves as a natural negative control, because this species lacks canonical RNAi machinery (Drinnenberg et al. 2009): even the few matches it carries could not produce canonical RNAi silencing.

### Designability: the hazard is avoidable in some genes and unavoidable in others

Hazard at the gene level does not imply that every construct against that gene is hazardous. To test whether hazard can be designed around, a window of 250 consecutive 21-mer start positions, spanning 270 nt of transcript and thus a realistic construct length, was slid along each gene in steps of one start position, and each window was scored for the presence of any hazardous 21-mer (Table 3; Figure 1). The results divide the target genes into three classes. CYP51C is entirely clean: all 1,285 possible windows contain zero hazardous 21-mers. CYP51A (60.0% clean windows), CHS1 (35.1%) and CYP51B (26.6%) are designable: hazard is concentrated in defined regions, and a construct placed in a clean window avoids it entirely. TUB2b is not designable: zero of 1,379 possible windows are clean: every 270-nt window (250 consecutive 21-mer start positions) against *F. graminearum* β-tubulin contains at least one perfect 21-nt match to a beneficial fungus, and the worst window contains 100. One qualification applies to the construct-length interpretation: a literal 250-nt dsRNA spans only 230 k-mer start positions, and because a shorter window can be clean where a longer window is not, the TUB2b result demonstrates the hazard for windows of 250 k-mer starts, while whether any literal 250-nt window of TUB2b is hazard-free is not directly established by this analysis and would require re-analysis with a 230-start window. The positional landscapes (Figure 1) show the same structure spatially: hazardous start positions comprise 0.3% of 21-mer starts in CYP51A, 5.8% in CYP51B, 0.0% in CYP51C, 6.7% in CHS1 and 19.3% in TUB2b, in each case concentrated in discrete peaks corresponding to conserved blocks of the orthologous genes rather than scattered uniformly. The design implication is immediate for the flagship construct: because 125 of the 129 hits in CYP51-concat derive from CYP51B, a redesigned CYP3RNA that repositions, rather than removes, the hazardous CYP51B window to one of the gene’s hazard-free regions would be expected to lower the off-target load while retaining efficacy against *Fusarium* at a fraction of the off-target sequence load.

## Discussion

This study set out to answer a narrow question: how often does a dsRNA designed anywhere within a canonical antifungal target gene of *Fusarium graminearum* contain a 21-nt stretch perfectly matching a transcript in a beneficial or non-target fungus? The answer, across the constructs screened, is that such matches are common. Eleven of twelve constructs carried off-target sequence hazard against a seven-species non-target panel; hazard reached 580 perfect siRNA-length matches in TUB2b, for which no hazard-free window of 250 consecutive 21-mer start positions (spanning 270 nt of transcript) exists; and the matches landed on orthologues of the targeted essential genes in the very fungi deployed in agriculture as biocontrol agents. Equally important is what the screen does not show. It is a hazard screen, not a risk assessment. A perfect 21-nt match demonstrates that a potential silencing trigger is present in the sequence; whether that trigger ever produces an effect in the field depends on environmental persistence of the dsRNA, on exposure of the non-target organism, on cellular uptake, and on a functioning silencing pathway in the recipient. None of these downstream steps was tested here, and the numbers in Tables 2 and 3 must not be read as predictions of harm.

The strongest version of the reviewer objection is uptake, and it deserves a direct answer. Qiao et al. (2021) showed that the efficiency with which fungi internalise environmental dsRNA varies dramatically among species and predicts SIGS efficacy: *Botrytis cinerea*, *Sclerotinia sclerotiorum*, *Verticillium dahliae* and *F. graminearum* take up dsRNA efficiently, whereas several other fungi do not. Critically for the present results, the beneficial *Trichoderma virens* showed weak uptake. If uptake is the rate-limiting barrier to off-target silencing (Šečić and Kogel 2021), then the 133 perfect TUB2b matches in *T. virens* may never become effects. Three considerations nonetheless argue against dismissing the hazard. First, uptake competence is known for only a handful of species; it is unknown for *R. irregularis*, unmeasured for most of the strains actually deployed in commercial products, and may differ between laboratory conditions and the field. Second, weak uptake is not no uptake, and exposure under a repeated-spray regime is not a single-dose event. Third, the same uptake literature cuts the other way for the efficacy claim itself: *Trichoderma* and *Fusarium* are both Hypocreales, and if future work shows efficient uptake in deployed *Trichoderma* strains, the perfect matches reported here become the highest-priority candidates for experimental testing. The correct reading of Qiao et al. (2021) is therefore not that the hazard is benign, but that hazard and risk are different quantities and this study measured the first.

Exposure is similarly double-edged. Naked dsRNA is short-lived in soil, degrading with half-lives of hours to a few days in agricultural soils, with adsorption to soil particles reducing bioavailability even as it transiently protects the molecule from nucleases (Dubelman et al. 2014; Parker et al. 2019). A soil-dwelling mycorrhizal fungus would therefore encounter sprayed dsRNA only briefly and at declining concentration. This reassurance weakens once delivery is engineered: formulation and nanocarrier technologies are designed not merely to prolong persistence but to promote plant uptake, apoplastic and symplastic transport, and root-directed delivery, so that root-applied or systemically translocated dsRNA can accumulate in root tissue (Sede et al. 2026), and under such regimes the exposure of root-associated beneficials, including arbuscular mycorrhizal fungi and root endophytes, to intact dsRNA is expected to rise rather than remain negligible. On foliage, however, dsRNA can remain biologically active for days after application (San Miguel and Scott 2016), and formulation technologies are being developed precisely to extend that persistence (Mitter et al. 2017). The realistic exposure scenario for phyllosphere and epiphytic fungi (including the *Trichoderma* and *Beauveria* strains sprayed onto the same crop surfaces) is thus days of direct contact, not hours. The insecticide literature again supplies the calibration: where 21-nt matches have been found in non-target arthropods and followed up experimentally, effects have generally been absent at field-realistic doses (Krishnan et al. 2021; Bulgarella et al. 2025). Match is not effect; but in every one of those precedents the match was found first and the experiment followed, which is exactly the sequence of events this screen is designed to initiate for fungi.

The orthologue pattern in Section 3.2 is biologically coherent, and its coherence is the finding. Perfect 21-nt matches between species are vanishingly unlikely to arise by chance at the densities observed; they arise because CYP51, chitin synthase and β-tubulin are essential genes under strong purifying selection whose orthologues retain long identical stretches among related fungi. The target genes of antifungal dsRNA design were chosen for exactly this property (essentiality and conservation make them robust fungicidal targets), and the same property guarantees that their off-target hazard concentrates in related fungi carrying the same essential genes. Cross-silencing driven by shared identical stretches is not hypothetical in this system: the CYP3RNA construct works by silencing all three CYP51 paralogues simultaneously through shared sequence (Koch et al. 2016), and longer constructs generate broader siRNA pools with greater cross-silencing potential (Höfle et al. 2020). The screen simply shows that the same mechanism, applied across a species boundary, reaches the biocontrol agents next door. The phylogenetic concentration of hits in Hypocreales matches the relatedness–spectrum rule established for insecticidal dsRNAs (Bachman et al. 2013) and converts that rule from a reassurance into a warning for the antifungal case.

For construct design, the results are cautiously encouraging. Off-target hazard is neither uniform nor immutable: it is a measurable, mappable property of a sequence, and in three of four hazardous genes examined in the designability analysis it can be eliminated entirely by placing the construct in a clean window (Table 3; Figure 1). CYP51C emerges as a naturally clean target, and the CYP3RNA redesign logic is explicit: 125 of its 129 modelled hits derive from the CYP51B component, so repositioning that window to a hazard-free region of CYP51B should lower the off-target load; because Koch et al. (2013) found that silencing all three CYP51 paralogues simultaneously was more efficient than targeting any single paralogue, however, removing CYP51B outright could itself cost efficacy, making repositioning within the gene the more defensible redesign. Gene selection and window placement are, in effect, specificity tools that cost nothing beyond a screen of the kind reported here, and tools for siRNA-directed construct design already exist to operationalise this (Lück et al. 2019; Cedden et al. 2025). The exception is as informative as the rule: TUB2b, with zero clean windows and a worst window of 100 hazardous 21-mers, should be treated as an un-designable target against this non-target panel, and constructs against conserved housekeeping targets of that class should attract the closest regulatory scrutiny.

A comparison with the registered sprayable precedent makes the gap concrete. In the United States Environmental Protection Agency’s assessment of ledprona, bioinformatic analysis found 417 twenty-one-nucleotide sequence matches to the target Colorado potato beetle transcript against a maximum of three in any single non-target species (US EPA 2023), a specificity margin of more than two orders of magnitude that supported registration. The present results show that nothing comparable can be assumed for antifungal constructs: the TUB2b construct carries 580 perfect 21-mer matches across six non-target species (577 of them in the five beneficial Hypocreales, with the remaining three in *R. irregularis*), and even the CYP51-concat concatenation carries 129. The difference is not in the quality of the constructs but in the biology of the target–non-target relationship (beetle versus earthworm is a long evolutionary distance, *Fusarium* versus *Trichoderma* is not) and in the composition of the screening panel. Ledprona’s margin was measured against the guideline non-target species; the antifungal hazard reported here was only visible once beneficial fungal transcriptomes were added to the panel.

For the emerging regulatory frameworks, the study identifies a blind spot with a straightforward remedy. The De Neef et al. (2026) framework prescribes systematic screening of dsRNA sequences against non-target transcriptomes using the ≥21-nt perfect-match and ∼80% identity criteria, and its human-health counterpart has already been applied to a registered product (Devisetty et al. 2025). But the non-target organism sets against which such screens are run are inherited from exposure-driven chemical pesticide assessment and contain no fungus (Romeis and Widmer 2020; Dietz-Pfeilstetter et al. 2021). Had the CYP3RNA design been screened under current practice, its 125 CYP51B-derived matches to *Trichoderma*, *Beauveria* and *Metarhizium* would not have been detected, because none of those transcriptomes would have been in the panel. Adding the deployed beneficial fungi, and as AMF testing matures *R. irregularis* as well (Mallmann et al. 2018; Roshanfekrrad et al. 2025; Papadopoulos et al. 2025), to the standard screen is inexpensive and closes the gap. The biocontrol-compatibility implication deserves emphasis in its own right: *Trichoderma* and *Metarhizium*-based products are co-deployed on the same crops that future antifungal dsRNA sprays would treat, and *M. brunneum*, represented by its sequenced reference-genome strain ARSEF 3297, which carried 126 perfect TUB2b matches (the commercially deployed strains of this species, such as F52/BIPESCO 5, are approved as active substances in the EU and US), belongs to the commercially deployed *Metarhizium* biocontrol agents, a class of strain-level products approved as active substances in the EU and US. A hazard screen that excludes these species cannot certify compatibility between the two fastest-growing classes of biological crop protection.

The clean results deserve as much weight as the hazardous ones. *R. irregularis* carried zero matches against the CYP51-concat concatenation and at most ten against any construct; given the central role of arbuscular mycorrhizal fungi in nutrient uptake and ecosystem functioning (Powell and Rillig 2018), the absence of perfect-match hazard in the model AMF is a genuinely positive finding for the mycorrhizal protection goal, pending uptake data. *S. cerevisiae*, which lacks canonical RNAi machinery (Drinnenberg et al. 2009), was nearly clean and in any case could not convert matches into silencing, serving as the screen’s natural negative control.

Six limitations bound the interpretation. First, whole target transcripts were screened rather than the actual published construct fragments; the numbers describe the hazard of designing anywhere within each gene, and a corpus audit of the published constructs remains to be done. Second, a single target pathogen was examined; extension to *Botrytis*, *Sclerotinia*, *Zymoseptoria* and *Magnaporthe* targets is required before generalising across antifungal pipelines. Third, the non-target panel comprised seven species, with no host plant and no pollinator outgroup such as *Apis mellifera*. Fourth, only the perfect 21-nt criterion was implemented; the ∼80% overall-identity criterion requires alignment, and mismatch-tolerant silencing below the perfect-match threshold is documented experimentally (Jackson et al. 2003; Kulkarni et al. 2006; Chen et al. 2021), so the screen understates total off-target sequence space. Conversely, the screen overstates the number of matches likely to be functional, because siRNA production from a long dsRNA is non-uniform, and not every theoretical 21-mer is diced into abundant siRNAs (Koch et al. 2016; Höfle et al. 2020); effective silencing further requires that the homologous stretch be accessible in the off-target transcript under physiological conditions, so the perfect-match counts bound the sequence-level hazard from above as well as from below. Fifth, the study measured hazard only: no exposure modelling and no uptake data were generated, and the weak dsRNA uptake reported for *T. virens* (Qiao et al. 2021) means that at least some of the hazard reported here may not translate into risk. Sixth, annotation quality varies across the reference assemblies; several hits in *B. bassiana* and *T. virens* land on transcripts annotated only as “hypothetical protein”, almost certainly the unannotated orthologues but not verifiably so without further analysis.

These limitations define the future work directly. The immediate priority is the corpus audit: screening the actual published construct sequences, beginning with CYP3RNA, against this and larger non-target panels. The panel should be expanded to additional target pathogens (the comparison species already staged in the reference panel, *Botrytis cinerea*, *Sclerotinia sclerotiorum*, *Verticillium dahliae* and *Zymoseptoria tritici*, together with *Magnaporthe*), to root endophytes such as *Serendipita indica*, to host plants, and to pollinator outgroups such as *Apis mellifera*; the 80% overall-identity criterion should be implemented to complete the two-tier screen of De Neef et al. (2026); the positional hazard map should be weighted by predicted siRNA yield and target-site accessibility, using established siRNA-design tools (Lück et al. 2019), so that experimental effort is directed at matches that are both abundantly diced and accessible in the recipient transcript; and the hits prioritised here, above all the β-tubulin and CYP51B matches in deployed *Trichoderma* and *Metarhizium* strains, should be tested experimentally for uptake and silencing in the recipient fungi, because only that experiment can convert the hazard map of this study into a risk estimate. Priority for those experiments should follow the joint gradient of hazard density and uptake competence: the Hypocreales biocontrol agents combine the heaviest match load with the closest relatedness to the target pathogen, and it is their response to field-formulated dsRNA that will determine whether the sequence-level hazard documented here is a manageable design constraint or a genuine obstacle to antifungal SIGS.

An exact, reproducible in-silico screen shows that perfect siRNA-length matches to beneficial fungi are common in canonical antifungal dsRNA targets of *Fusarium graminearum*: eleven of twelve constructs carried off-target sequence hazard, concentrated in the Hypocreales relatives of the target pathogen and landing on orthologues of the targeted essential genes, with β-tubulin carrying 580 matches and no hazard-free window of 250 consecutive 21-mer start positions (spanning 270 nt of transcript). The hazard is real but designable: CYP51C is naturally clean, most hazard in other genes is avoidable by window placement, and the flagship CYP3RNA design’s off-target load is almost entirely attributable to one repositionable component. It is also invisible to current regulatory practice because no non-target test battery or bioinformatic screen includes a fungus. These are hazard findings, not risk findings; converting them into risk estimates requires exposure, uptake and silencing data that do not yet exist for beneficial fungi. Generating those data, and adding beneficial fungi to routine off-target screening, should precede the registration of antifungal dsRNA sprays.

## Acknowledgements

We thank Karl-Heinz Kogel (Justus Liebig University Giessen) for helpful comments on an earlier version of this manuscript.

## Statements and Declarations

## Funding

The authors did not receive support from any organization for the submitted work.

### Ethics approval

Not applicable.

### Use of generative AI and AI-assisted technologies

During the preparation of this manuscript, the authors used generative artificial intelligence (large language model) tools to assist with language editing, with formatting the manuscript to the journal style, and with checking the bibliographic accuracy of the references. No generative AI tool was used to design the study, to generate, analyse or interpret the data, or to produce the scientific conclusions; the in-silico off-target screen was performed with the authors’ own Python code. The authors reviewed and edited all AI-assisted content and take full responsibility for the content of this publication.

### Data availability

All reference transcriptomes analysed in this study are publicly available from Ensembl Fungi (downloaded 3 August 2026; Ensembl Fungi release 63). The analysis code (the k-mer screening engine kmerlib.py and the companion fetch, screen and landscape scripts), the reference-panel manifest and all result files are openly available at Harvard Dataverse: https://doi.org/10.7910/DVN/ZDLO7W (CC0 1.0). Analyses were run in Python 3 using numpy, Biopython and matplotlib, as documented in the deposited README. The method is fully described in the Materials and methods section.

### Author contributions

Conceptualization: Muhammad Wasif; Methodology: Muhammad Wasif; Software: Muhammad Wasif, Muhammad Ahsan Khan; Validation: Huma Saleem, Gábor Tarcali; Formal analysis: Muhammad Ahsan Khan; Investigation: Huma Saleem; Data curation: Muhammad Ahsan Khan; Writing – original draft preparation: Muhammad Wasif; Writing – review and editing: Muhammad Wasif, Gábor Tarcali, Muhammad Ahsan Khan, Huma Saleem, Muhammad Abubakkar Azmat; Visualization: Muhammad Wasif; Supervision: Gábor Tarcali, Muhammad Ahsan Khan; Project administration: Gábor Tarcali. All authors read and approved the final manuscript.

